# T3SS translocon induces pyroptosis by direct interaction with NLRC4/NAIP inflammasome

**DOI:** 10.1101/2024.07.11.603062

**Authors:** Yan Zhao, Hanshuo Zhu, Jinqian Li, Hang Xu, Li Sun

**Affiliations:** CAS and Shandong Province Key Laboratory of Experimental Marine Biology, Institute of Oceanology; CAS Center for Ocean Mega-Science, Chinese Academy of Sciences, Qingdao, China; Laboratory for Marine Biology and Biotechnology, Qingdao Marine Science and Technology Center, Qingdao, China; College of Marine Sciences, University of Chinese Academy of Sciences, Qingdao, China; Tsinghua University-Peking University Joint Center for Life Sciences, School of Basic Medical Sciences, Tsinghua University, Beijing 100084, China; NHC Key Laboratory of Tropical Disease Control, School of Tropical Medicine, Hainan Medical University, Haikou, Hainan, 571199, China

**Keywords:** T3SS, translocon, NLRC4, pathogen-host interaction

## Abstract

Type III secretion system (T3SS) is a virulence apparatus existing in many bacterial pathogens. Structurally, T3SS consists of the base, needle, tip, and translocon. The NLRC4 inflammasome is the major receptor for T3SS needle and basal rod proteins. Whether other T3SS components are recognized by NLRC4 is unclear. In this study, using *Edwardsiella tarda* as a model intracellular pathogen, we examined T3SS−inflammasome interaction and its effect on cell death. *E. tarda* induced pyroptosis in a manner that required the bacterial translocon and the host inflammasome proteins of NLRC4, NLRP3, ASC, and caspase 1/4. The translocon protein EseB triggered NLRC4/NAIP-mediated pyroptosis by binding NAIP via its C-terminal region, particularly the terminal 6 residues (T6R). EseB homologs exist widely in T3SS-positive bacteria and share high identities in T6R. Like *E. tarda* EseB, all of the representatives of the EseB homologs exhibited T6R-dependent NLRC4 activation ability. Together these results revealed the function and molecular mechanism of EseB to induce host cell pyroptosis and suggested a highly conserved inflammasome-activation mechanism of T3SS translocon in bacterial pathogens.

## Introduction

The host innate immune system responds to pathogen-associated molecular patterns (PAMPs) and damage-associated molecular patterns (DAMPs) via multiple pattern recognition receptors (PRRs). Inflammasomes are a group of cytoplasmic PRRs that detect intracellular pathogens or disruptions in cellular homeostasis [1]. NLRP1, NLRP3, NLRC4, AIM2 and Pyrin are well-established PRRs that always combine with the adaptor protein ASC to form canonical inflammasomes, which activate the effector protein caspase-1 (Casp1), leading to the processing and release of interleukin (IL) -1β and IL-18 [2]. Casp1 can also cleave and activate gasdermin (GSDM) D, which subsequently forms channels in the plasma membrane, eventually leading to a type of lytic programmed cell death called pyroptosis [3, 4]. In the non-canonical pathway, Casp4/5 (in humans) and Casp11 (in mice) are activated by bacterial lipopolysaccharide (LPS) and trigger GSDMD-mediated pyroptosis [4, 5]. Of these inflammasomes, NLRC4 responds to Gram-negative bacterial ligands, primarily flagellin and components of the type III secretion system (T3SS) apparatus, in a manner that requires an upstream immune sensor protein called NLR-family apoptosis inhibitory protein (NAIP), which interacts directly with the PAMPs [6-9]. Mice possess several NAIPs, each of which detects specific bacterial ligands, while humans possess only one functional NAIP (hNAIP) that is capable of broadly recognizing multiple bacterial ligands [7-10].

*Edwardsiella tarda* belongs to the Enterobacteriaceae family. It is an intracellular pathogen and can survive and replicate in host immune cells, such as macrophages [11, 12]. *E. tarda* has a broad range of hosts, including fish and humans [13, 14]. In humans, *E. tarda* has been reported to cause gastrointestinal diseases and systemic infection that can be lethal [15, 16]. *E. tarda* possesses T3SS and uses it to modulate the host immune systems [13, 16]. T3SS functions as an injectisome that delivers bacterial effector proteins into host cells. The T3SS apparatus consists of three distinct parts— the extracellular segment, the basal body, and the cytoplasmic components [17]. The extracellular part comprises the needle, tip, and translocon, which spans the host cell membrane [18]. In *E. tarda*, the translocon complex is formed by EseB, EseC, and EseD [19]. Genetically, the *eseB*, *escA*, *eseC*, and *eseD* genes clustered tandem in the same operon. In function, EscA acts as a specific chaperone for EseC [20]. The translocon is known to be essential for the pathogenesis of *E. tarda* [19-21], but the mechanism, in particular that associated with inflammasome activation and pyroptosis, remains to be explored.

In this study, using *E. tarda* as an intracellular pathogen model and human macrophages as the host cells, we investigated the function and the working mechanism of the T3SS translocon in pathogen-host interaction. We found that *E. tarda* induced GSDMD-dependent pyroptosis involving both canonical and non-canonical inflammasomes, and that the translocon proteins were indispensable for *E. tarda* cytotoxicity. We examined the role and mechanism of EseB in host interaction and uncovered the key structure of EseB that was essential for binding and activating the NLRC4/NAIP inflammasome. Furthermore, we identified EseB homologs in a broad range of bacteria and demonstrated that NLRC4/NAIP-interaction and activation was probably a conserved function of the EseB homologs in T3SS-positive bacterial pathogens. These results added new insight into the working mechanism of EseB and highlighted the important role of the translocon in bacteria-host interaction.

## Material and Methods

### Cells and cell culture

HEK293T and THP-1 cells were purchased from American type culture collection, ATCC (Manassas, USA) and Cell Resource Center, IBMS, CAMS/PUMC (Beijing, China), respectively. The cells were maintained at 37 °C in a 5% CO_2_ humidified incubator. HEK293T cells were cultured in DEME (C11995500, Gibco) supplemented with 10% (v/v) FBS (10099-141C, Gibco), 1% penicillin, and streptomycin (SV30010, HyClone). THP-1 cells were cultured in complete RPMI 1640 medium composed of RPMI 1640 (C22400500, Gibco) medium supplemented with 10% (v/v) FBS and 1% penicillin and streptomycin. THP1-Null (control), THP1-Casp1-KD (*Casp1* knockdown), and THP1-NLRP3-KD (*Nlrp3* knockdown) were obtained from InvivoGen (Hong Kong, China) and maintained as instructed by the manufacturer.

### Gene knockout and knockdown

THP-1 cells with gene knockout were generated using the CRISPR-Cas9 system as de scribed previously [22, 23]. Briefly, the sgRNAs targeting Aim2 (5ʹ-TTCACGTTTGA GACCCAAGA-3ʹ), Casp4 (5ʹ-TGGTGTTTTGGATAACTTGG-3ʹ), and NLRC4 (5ʹ-CCA CTACCACTGAGTGCCTG-3’) were used for lentiviral constructs. The cells were treat ed with the lentiviral constructs and selected with puromycin. After selection, the gene knockout cells derived from single cells were further verified by PCR and sequence analysis. For *NAIP* gene knockdown via short hairpin RNA (shRNA), the oligo targeting *NAIP* (5’-CCGGGCCGTGGTGAACTTTGTGAATCTCGAGATTCACAAAGTTCACCACGGCTTTTTG-3’ and 5’-AATTCAAAAAGCCGTGGTGAACTTTGTGAATCTCGAGATTCACAAAGTTCACCACGGC-3’) was cloned into pLKO.1 puro (8453, Addgene), which was then used for lentiviral construct as above. The pLKO.1 scramble (1864, Addgene) was used for creating the negative control lentiviral construct. THP-1 cells wer e treated with the lentiviral constructs and selected as above. The knockdown efficiency was verified by RT-PCR with primers F (5’-GGCCAAACTGATCATCCAGC-3’) and R (5’-TGGCATGTTGTCCAGTGCTT-3’).

### Bacterial strains and culturing

The *E. tarda* mutants with markerless in-frame deletion of *eseB*, *escA*, *eseC*, *eseD*, *eseB*-*eseD* and *fliC1/2* (Δ*eseB*, Δ*escA*, Δ*eseC*, Δ*eseD*, Δ*eseB-D* and Δ*fliC*, respectively) were constructed as reported previously [24, 25]. Briefly, the fragments upstream and downstream of the target gene were amplified by overlapping PCR and inserted into the suicide plasmid pDM4. The recombinant plasmids were introduced into *E. tarda*, and mutant strains were generated by a two-step homologous recombination. The deletion of the target gene was confirmed by PCR and sequence analysis of the PCR products. The information of primers and the mutants is shown in Table S1. *E. tarda* and its mutants were grown in Luria–Bertani (LB) medium supplemented with 20 μg/mL polymycin B (P8350, Solarbio) at 28 °C.

### Purification of recombinant proteins

The coding sequences of EscA, EseB, EseC, and EseD were amplified by PCR from the genome of *E. tarda*. All PCR products were cloned into the plasmid pET-28a (Novagen, 69864). *E. coli* BL21(DE3) (CD601, TransGen Biotech) was transformed with each of the recombinant plasmids, and the transformant was grown in LB medium at 37°C until OD_600_ 0.6. Isopropyl-β-D-thiogalactopyranoside (I8070, Solarbio) (0.2 mM) was added to the bacterial culture, and the culture was continued overnight at 16 °C. Bacteria were collected and lysed in Buffer A (20 mM Tris-HCl pH 8.0, 300 mM NaCl, and 10 mM imidazole). The recombinant proteins with His-tag were purified with Ni-NTA Agarose (30210, Qiagen). The proteins loaded onto the Ni-NTA column were washed with 60 column volumes of the Buffer B (20 mM Tris-HCl pH 8.0, 300 mM NaCl, 40 mM imidazole, and 0.1% Triton X114), and then with 80 column volumes of Buffer C (20 mM Tris-HCl pH 8.0, 300 mM NaCl, and 40 mM imidazole). The proteins were finally eluted with Buffer D (20 mM Tris-HCl pH 8.0, 300 mM NaCl, and 250 mM imidazole) and dialyzed against Buffer E (20 mM Tris-HCl pH 8.0, and 150 mM NaCl). The purified proteins were subjected to SDS–PAGE. Protein concentrations were determined with the BCA Protein Assay Kit (P0010, Beyotime) according to the manufacturer’s instruction.

### *E. tarda* infection in THP-1 cells

THP-1 cells were differentiated into macrophages with PMA overnight [26]. *E. tarda* variants were cultured in LB medium with 20 μg/mL polymycin B at 28 °C until OD_600_ 0.8. The bacteria were washed with PBS twice and then mixed with the differentiated THP-1 cells at MOI=10. The mixture was centrifuged at 800*g* for 8 min and incubated at 30 °C for 1h in a 5% CO_2_ humidified incubator. To kill the extracellular bacteria, gentamycin (500 μg/mL) was added to the cells, followed by incubation for 0.5 h. The culture medium was replaced with fresh medium containing 40 μg/mL gentamycin. To prevent bacterial entry into cells, the cells were treated with 50 μM cytochalasin B (ab143482, Abcam) or 10 μM cytochalasin D (PHZ1063, Invitrogen) for 0.5 h prior to infection.

### Protein electroporation

THP-1 cells were cultured in complete RPMI 1640 medium to a density of ∼1×10^6^ cells/mL. The cells were washed with precooled cytoporation medium T (47-0002, BTXpress) twice and resuspended in medium T to 5×10^6^ cells/mL. Protein electroporation was performed using a Gemini X2 electroporator (45-2006, BTX) as follows. The protein (2 μg) was added to 1×10^6^ cells in 200 μl medium T. Electroporation was then performed at the setting of 300 V, 10 ms, and 1 pulse. The cells were transferred into 1 mL pre-warmed OPTI-MEM medium (31985070, Gibco) and incubated for 1 h. Then both cell lysates and supernatants were collected as described previously [22, 26] for immunoblotting. Cell death was determined with CytoTox 96^®^ Non-Radioactive Cytotoxicity Assay kit (G1780, Promega).

### NLRC4 inflammasome reconstitution in HEK293T cells

The coding sequences of human NLRC4, NAIP, Casp1, and proIL-1β were cloned from PMA-differential THP-1 cells and inserted into pCS2Flag (16331, Addgene)-based expression vectors with different tags. The coding sequences of EseB homologs from various bacteria (GenBank accession numbers shown in Table S2) were synthesized by Sangong Biotech (Shanghai, China). For NLRC4 inflammasome reconstitution, HEK293T cells were seeded into 6-well plates overnight and then transfected with the indicated combination of plasmids (2 μg, 100 ng, 100 ng, 25 ng, and 100 ng for the plasmids expressing proIL-1β, NLRC4, NAIP, Casp1, and EseB, respectively) using Lipofectamine 3000 (L3000015, Invitrogen). At 24 h post transfection, the cells were lysed using RIPA buffer containing protease inhibitors. The cell lysates were analyzed by immunoblotting as described below.

### Immunoblotting and immunoprecipitation

Immunoblot was performed as reported previously [26] with the following antibodies: Caspase-1 antibody (2225S, Cell Signaling Technology), GSDMD antibody (96458S, Cell Signaling Technology), IL-1β rabbit mAb (12703S, Cell Signaling Technology), Caspase-4 mAb (42264T, Cell Signaling Technology), anti-6×His tag mAb (ab213204, Abcam), flag-tag rabbit mAb (AE063, ABclonal), mouse anti HA-Tag mAb (AE008, ABclonal), mouse anti Myc-tag mAb (AE010, ABclonal), β-actin mouse mAb (AC004, ABclonal), HRP goat anti-mouse IgG (H+L) (AS003, ABclonal), goat anti-rabbit IgG H&L (HRP) (ab97051, Abcam). For immunoprecipitation, HEK293T cells transfected with the indicated plasmids were lysed in IP lysis buffer (50 mM Tris-HCl pH 7.6, 150 mM NaCl, 1% triton X-100, and 1×protease inhibitor cocktail), followed by centrifugation at 14,000*g* for 10 min to remove cell debris. The supernatants were mixed with equilibrated anti-FLAG® M2 magnetic beads (M8823, sigma) according to the manufacturer’s instructions.

### Data analysis and statistics

Data were analyzed primarily using the Prism 10 software (www.graphpad.com). Statistical analysis was conducted using Student’s *t*-test for comparing two sets of data and one-way ANOVA for comparing three or more sets of data. Significance was defined as \**p* < 0.05, \*\**p* < 0.01, \*\*\**p* < 0.001 and \*\*\*\**p* < 0.0001.

## Results

### 3.1 *E. tarda* triggers GSDMD-dependent pyroptosis in human macrophages

To examine whether *E. tarda* infection triggered cell death in human macrophages, differentiated THP-1 cells (dTHP-1 cells) were infected with *E. tarda*. The cells were found to undergo rapid cell death as indicated by LDH release (Figure S1A). However, cell death was almost completely blocked when the cells were pre-treated with cytochalasin B (CytoB) or cytochalasin D (CytoD), which inhibited bacterial entry into the cells (Figure S1A, B). Hence, it was intracellular *E. tarda*, rather than extracellular bacteria, that induced cell death in human macrophages. Further examination showed that *E. tarda*-infected dTHP-1 cells exhibited a swelling morphology, accompanied with IL-1β release, Casp1 activation, and GSDMD cleavage (Figure 1A-E). These observations indicated that *E. tarda* triggered pyroptosis in dTHP-1 cells. To examine whether and which inflammasomes were involved in this process, cells defective in various inflammasome pathways were employed. The results showed that following *E. tarda* infection for 2 h or 4 h, cell death and IL-1β release were significantly decreased in NLRC4 knockout (NLRC4-KO) cells, Casp4 knockout (Casp4-KO) cells, and NLRP3 knockdown (NLRP3-KD) cells (at 4 h post-infection), but not in Aim2 knockout (Aim2-KO) cells (Figure 1F-I). Cells with Casp1 knockdown (Casp1-KD), GSDMD knockout (GSDMD-KO), and ASC knockout (ASC -KO) all exhibited significantly decreased cell death and IL-1β release (Figure 1F-I). Taking together, these results indicated that intracellular *E. tarda* induced GSDMD-dependent pyroptosis in human macrophages in a manner that required NLRC4, NLRP3, ASC, Casp1, and Casp4.

**Figure 1.**
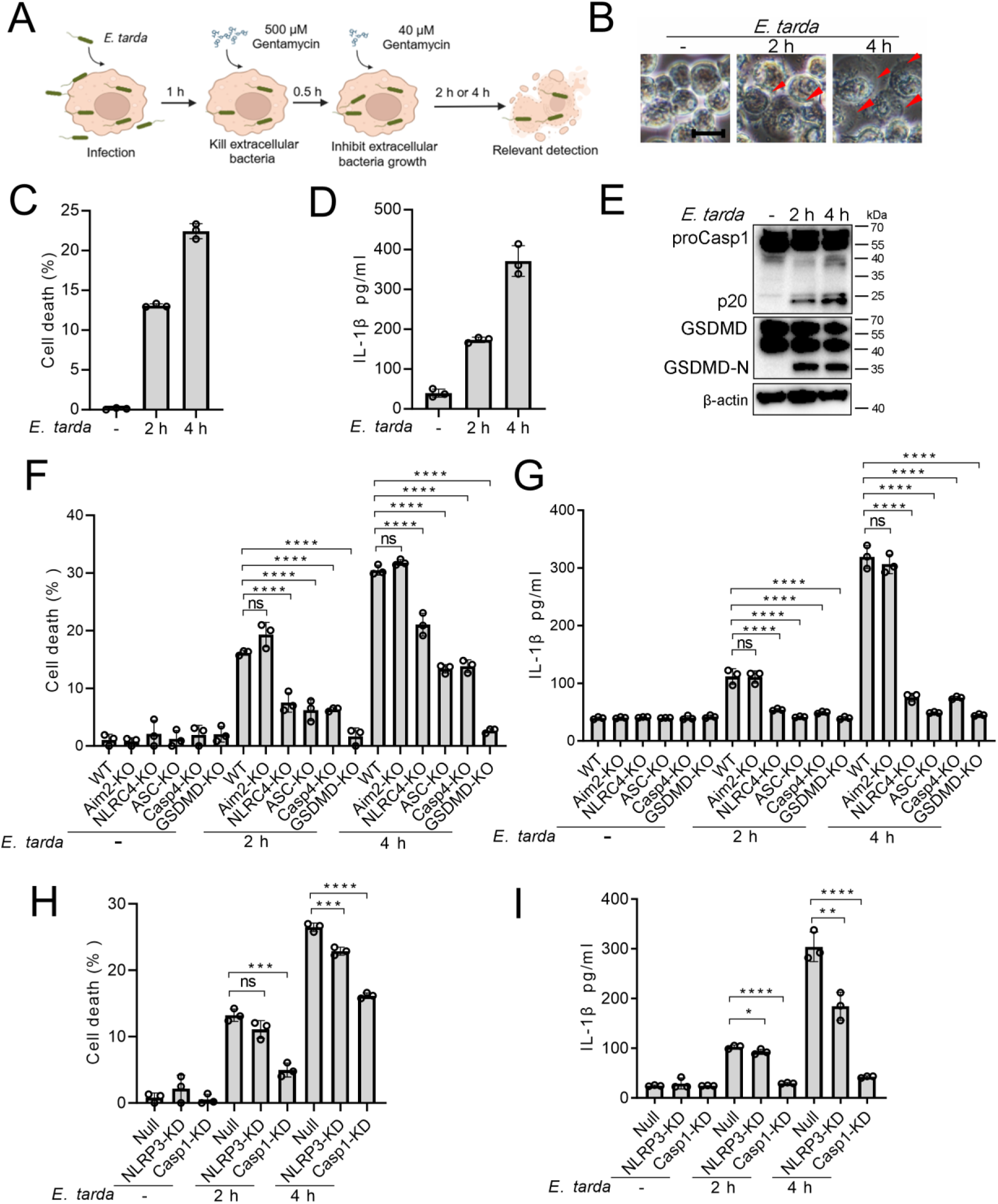
The ability of *Edwardsiella tarda* to induce pyroptosis in human macrophages. (**A**) The schematic of experimental design. (**B-E**) dTHP-1 cells were infected with *E. tarda* for the indicated hours and then subjected to microscopy (B), measurement of cell death (C), IL-1β release (D), and Western blot (E) using antibodies against Casp1, GSDMD, and β-actin (loading control). In (B), red arrowheads indicate pyroptotic cells; scale bar, 10 μm. (**F-I**) dTHP-cells in the form of wild type (WT), knockout (KO) variants (Aim2-KO, NLRC4-KO, ASC-KO, Casp4-KO, and GSDMD-KO), and knockdown (KD) variants (NLRP3-KD and Casp1-KD) were infected with or without *E. tarda* for 2 or 4 h, and then assessed for cell death (F, H) and IL-1β release (G, I). For panels C, D, and F-I, data were the means of triplicate assays and shown as means ±SD. ns, not significant, \*\*\**p*<0.001, \*\*\*\**p*<0.0001, one-way ANOVA with Dunnett’s multiple-comparison test.

### 3.2 The T3SS translocon is essential to *E. tarda-*induced pyroptosis

Similar to most intracellular Gram-negative bacterial pathogens, *E. tarda* possesses a T3SS system and uses it as a weapon against host immunity. In this system, EseB, EseC, and EseD form a translocon, with EscA acting as an EseC chaperone. To investigate the potential role of the translocon in pyroptosis, a series of *E. tarda* mutants were constructed, which bear the knockout of *eseB*, *escA*, *eseC*, or *eseD* (Δ*eseB*, Δ*escA*, Δ*eseC*, or Δ*eseD*, respectively), or the knockout of all of the four genes (Δ*eseB-D*) (Figure 2A). Compared with the wild type, these mutants showed no deficiency in host cell adhesion or entry (Figure 2B, C). However, Δ*eseB-D*, Δ*eseB*, Δ*eseC*, and Δ*eseD* were unable to induce host cell death, IL-1β release, Casp1 activation, or GSDMD cleavage following infection (Figure 2D-F). Casp4 activation was also absent in Δ*eseB-D*-infected cells (Figure S2). Δ*escA* was still able to induce pyroptosis, but the levels of cell death and IL-1β release were significantly lower than that induced by the wild type (Figure 2D-F). To examine whether flagellin was required for *E. tarda*-induced pyroptosis, the flagellin mutant, Δ*fliC*, was created. Compared with the wild type (WT), Δ*fliC* exhibited no significant change in host cell adhesion/entry, cell death induction, Casp1 activation, or GSDMD cleavage (Figure S3). These results indicated that the T3SS translocon proteins, rather than flagellin, were indispensable for *E. tarda*-induced pyroptosis.

**Figure 2.**
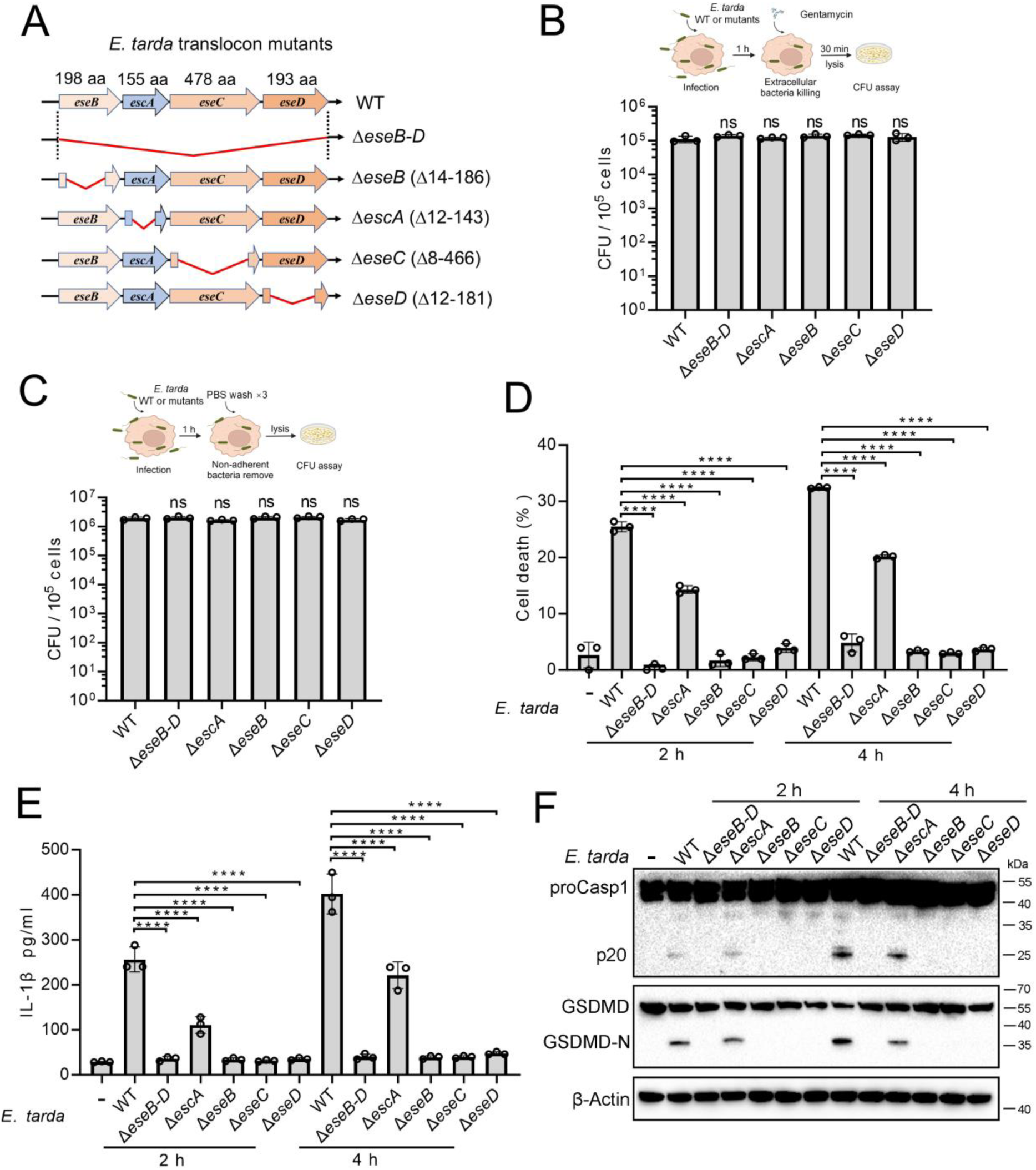
The importance of the translocon for *Edwardsiella tarda-*induced pyroptosis. (**A**) A diagram showing the in-frame deletion (red curved line) of *eseB-D*, *escA*, *eseB*, *eseC* and *eseD*. (**B, C**) dTHP-1 cells were infected with wild type (WT) or mutant *E. tarda* for 1h. The intracellular bacteria (B) and the total bacteria associated with the cells (i.e., both the cell-attached and the intracellular bacteria) (C) were determined by plate count. (**D-F**) dTHP-1 cells were treated with or without *E. tarda* variants for 2 or 4 h, and then subjected to cell death analysis (D), IL-1β release measurement (E), and immunoblot (F) using antibodies against Casp1, GSDMD, and β-actin (loading control). For panels B-E, data are the means of triplicate assays and shown as means ± SD. ns, not significant, \*\*\*\**p*<0.0001, one-way ANOVA with Dunnett’s multiple-comparison test.

### 3.3 Cytosolic EseB triggers pyroptosis in a NLRC4/NAIP-dependent manner

To examine the mechanism underlying the above observed essentialness of the translocon in *E. tarda*-induced cell death, the recombinant proteins of EscA, EseB, EseC, and EseD were prepared (Figure S4A). The effects of these proteins, both extracellular and intracellular, on THP-1 cells were determined. When present in the extracellular milieu, none of these proteins caused apparent change to the cell morphology (Figure 3A). However, when EseB was present in the cytoplasm of THP-1 cells, pyroptotic cell death, including activation of Casp1 and GSDMD, was observed (Figure 3A-C). To examine which inflammasome pathway was involved in this process, the effect of EseB was determined using Aim2/NLRC4/ASC/Casp4/GSDMD knockout cells and NLRP3/Casp1 knockdown cells. The results showed that defective in NLRC4, GSDMD, and Casp1, but not in AIM2, Casp4, or NLRP3, rendered the cells markedly immune to the death-inducing effect of EseB (Figure 3D, E). ASC knockout also significantly, though to a relatively modest extent, reduced EseB-induced cell death. Since NAIP is known to be involved in NLRC4 inflammasome activation, the effect of NAIP knockdown on EseB cytotoxicity was examined. The result showed that the NAIP-knockdown cells exhibited significantly decreased death following EseB treatment (Figure 3F, G). Collectively, these results indicated that cytosolic EseB, rather than extracellular EseB, induced pyroptosis via the NLRC4/NAIP inflammasome.

**Figure 3.**
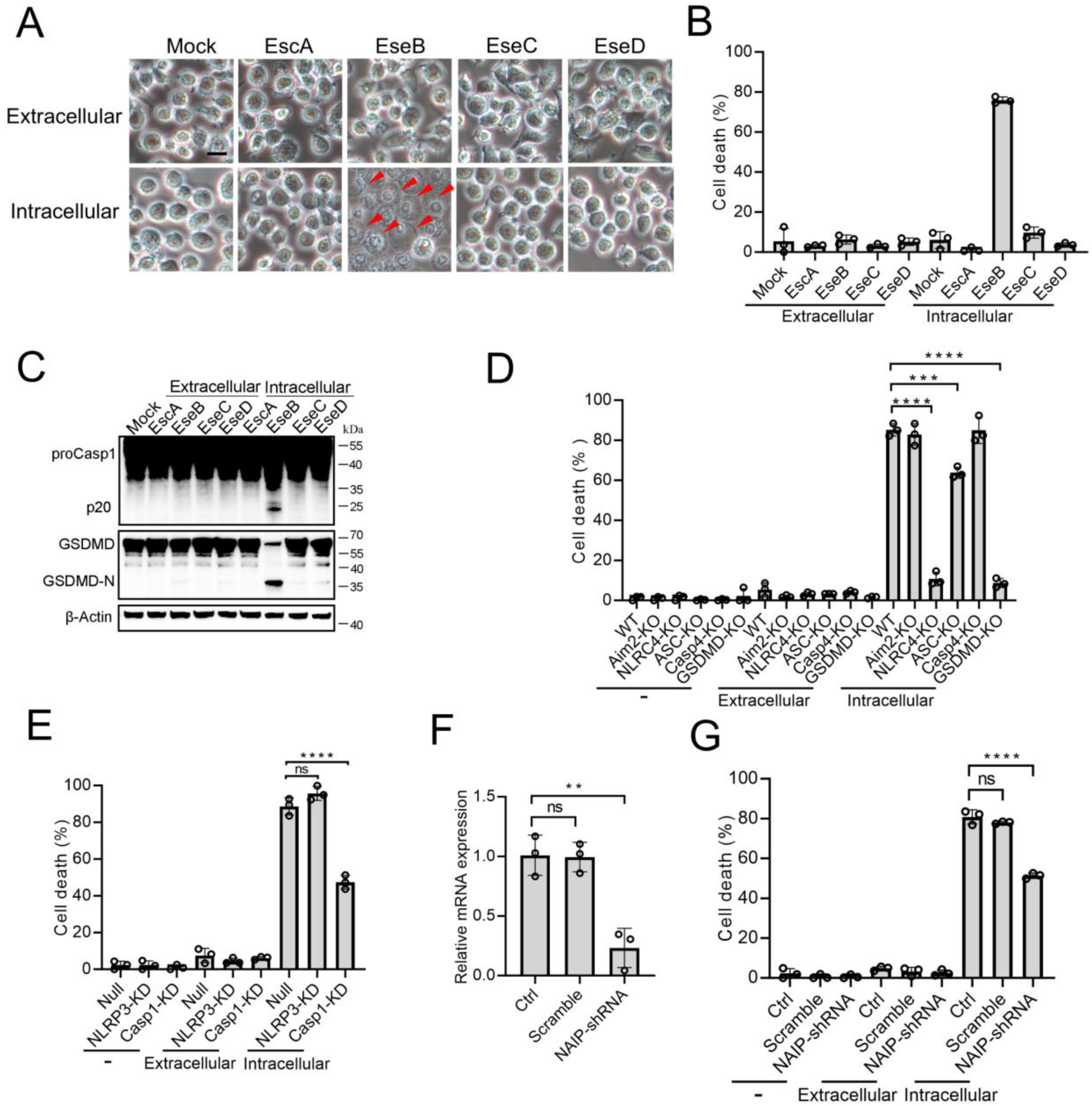
The pyroptotic effect of the translocon proteins and its dependence on the inflammasomes. (**A-C**) To determine the extracellular and intracellular effects of EscA, EseB, EseC, and EseD, each of the proteins was added into the culture medium of THP-1 cells (extracellular) or electroporated into THP-1 cells (intracellular). The control cells were mock treated with PBS. The cells were subjected to microscopy (A), cell death analysis (B), and immunoblot using antibodies against Casp1, GSDMD, and β-actin (loading control) (C). In (A), red arrowheads indicate pyroptotic cells; scale bar, 10 μm. (**D**) The wild type (WT) and knockout (KO) THP-1 cells were treated with or without extracellular and intracellular EseB as above and then examined for cell death. (**E**) The control THP-cells (Null) and the NLRP3/Casp1 knockdown (KD) THP-cells were treated with or without extracellular and intracellular EseB as above and then examined for cell death. (**F**) THP-1 cell treated with or without (Control, Ctrl) NAIP-targeting shRNA or scramble RNA (negative control RNA) were examined for NAIP expression by qRT-PCR. (**G**) THP-1 cells administered with or without NAIP-targeting or scramble RNA were treated or without extracellular and intracellular EseB as above and then examined for cell death. For panels B, and D-F, data are the means of triplicate assays and shown as means ± SD. ns, not significant, \*\*\**p*<0.001, \*\*\*\**p*<0.0001, one-way ANOVA with Dunnett’s multiple-comparison test.

### 3.4 The C-terminal region of EseB is vital to NAIP interaction and pyroptosis induction

To explore its mechanism to activate the NLRC4/NAIP pathway, EseB was first subjected to sequences analysis. The C-terminal (CT) region of EseB exhibits notable degrees of conservation, especially in the terminal 6 residues (T6R), with bacterial needle proteins known to activate NLRC4 (Figure 4A). Based on this observation, we divided EseB into the N-terminal (NT) (1-112 aa) and the CT (113-198 aa) regions (Figure 4B). To identify the sequence essential to EseB function, a series of EseB mutants were constructed that bear deletion of the terminal 4 residues (T4R) (EseBΔT4R) or the T6R (EseBΔT6R), or contain only the NT (EseB-NT) or CT (EseB-CT) region (Figure 4B, S4B). When introduced into THP-1 cells, EseB-CT induced cell death and Casp1/GSDMD activation, whereas all other EseB mutants failed to do so (Figure 4C-E). The EseB variants were further examined in 293T cells with reconstituted NAIP/NLRC4 inflammasome, in which NAIP/NLRC4 activation could be monitored by analyzing the maturation cleavage of IL-1β (Figure 4F). The results showed that in cells co-expressing EseB and all NLRC4 inflammasome components, massive IL-1β cleavage occurred (Figure 4G). Similar observations were made with cells co-expressing NAIP/NLRC4 inflammasome and EseB-CT (Figure S5A). Together these results demonstrated that EseB, via its CT region, activated the NAIP/NLRC4 inflammasome. Consistently, immunoprecipitation revealed that EseB, as well as EseB-CT, bound to NAIP, and this binding was not observed with either of the other EseB mutants (Figure 4H, I). EseB could not bind to NLRC4 directly (Figure 4H). In addition to EseB, the rod (EsaI), needle (EsaG), and flagellin (FliC1/2) of *E. tarda* were also examined for their ability to activate the NLRC4 inflammasome. For this purpose, these proteins were each co-expressed with NAIP/NLRC4 in 293T cells. Only the needle protein EsaG induced significant IL-1β cleavage (Figure S5B, C). This result indicated that EsaG, but not the rod protein or flagellin, activated NLRC4.

**Figure 4.**
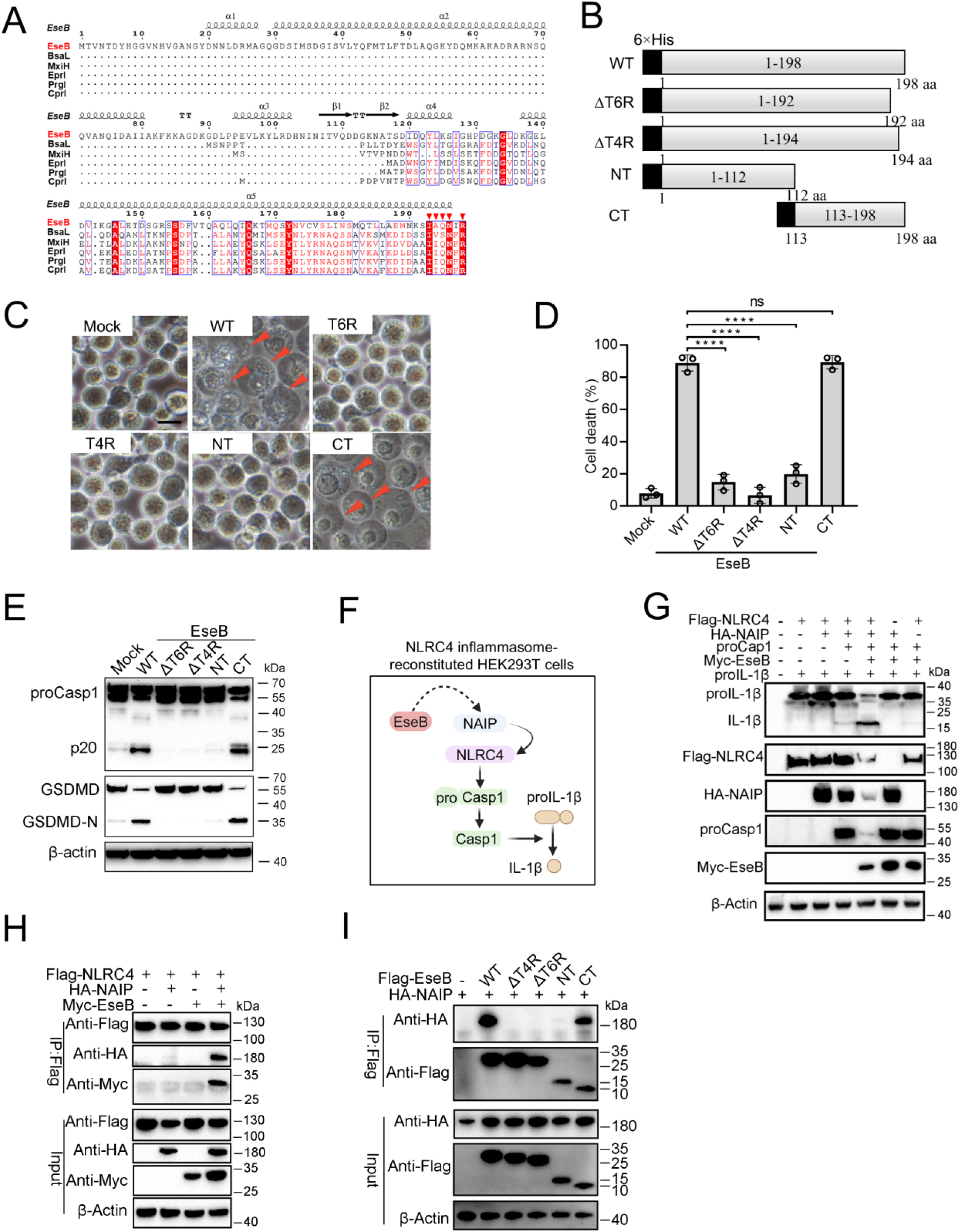
Identification of the functional important region in EseB. (**A**) Sequence alignment of EseB and T3SS needle proteins with NLRC4/NAIP-stimulating activity. (**B**) A diagram showing EseB wild type (WT) and truncates. (**C-E**) THP-1 cells were electroporated with or without (Mock) EseB WT or truncate. The cells were subjected to microscopy (C), cell death analysis (D), and immunoblot with antibodies against Casp1, GSDMD, and β-actin (loading control) (E). In (C), red arrowheads indicate pyroptotic cells; scale bar, 10 μm. For panel D, data are the means of triplicate assays and shown as means ± SD. ns, not significant, \*\*\*\**p*<0.0001, one-way ANOVA with Dunnett’s multiple-comparison test. (**F**) A diagram showing detection of the activating effect of EseB on NAIP/NLRC4 in NLRC4 inflammasome-reconstituted HEK293T cells by determining proIL-1β cleavage. (**G**) HEK293T cells were transfected with or without the indicated combination of vectors expressing Flag-tagged NLRC4, HA-tagged NAIP, Myc-tagged EseB, proCasp1, and proIL-1β for 24 h. The cells were subjected to immunoblot using antibodies against the tags or the proteins with β-actin as a loading control. (**H**) HEK293T cells were transfected with the indicated combination of vectors expressing Flag-tagged NLRC4, HA-tagged NAIP, and Myc-tagged EseB. The cells were subjected to immunoprecipitation (IP) using antibodies against the tags with β-actin as a loading control. (**I**) HEK293T cells were transfected with the indicated combination of vectors expressing HA-tagged NAIP and Flag-tagged EseB variants. IP was performed as above.

### 3.5 The NLRC4-activation capacity and mechanism of EseB are conserved in pathogenic bacteria

With the above results, we wondered whether the observed EseB function was unique to *E. tarda* or common in bacterial pathogens with T3SS. To answer this question, we searched and identified EseB homologs in diverse pathogenic bacteria. Among these EseB homologues, 20 were randomly selected, including that from *Salmonella* and *Chromobacterium*, for activity analysis (Table S2). The results showed that all of the 20 EseB homologs could activate the NLRC4/NAIP inflammasome in reconstituted 293T cells (Figure 5A). Phylogenetic analysis of these EseB homologues and *E. tarda* EseB showed that they fell into four groups (Figure S6). However, high levels of sequence identities are shared among these EseB at the T6R (Figure 5B). To examine whether, as observed with *E. tarda* EseB, the T6R was functionally important, five of the 20 EseB homologs were selected for mutation to remove the T6R or T7R. The resulting mutants, like *E. tarda* EseBΔT6R, no longer possessed the ability to activate the NLRC4 inflammasome and cause cell death (Figure 5C, D, S7). We further examined the NLRC4-activating potential of bacterial translocators with relatively low sequence identities (< 23%) with EseB. For this purpose, 14 key translocator proteins from 8 pathogenic bacteria were selected for the examination (Table S2). Three of these proteins, i.e., PcrV, SipC, and IpaC, possess terminal 5-7 residues that are similar to the T6R of EseB (Figure S8A). Subsequent study showed that only PcrV (from *Pseudomonas aeruginosa*) was able to activate the NLRC4 inflammasome (Figure S8B). When the terminal five residues (T5R) of PcrV were deleted, the resulting ΔT5R mutant lost the capacity to activate NLRC4 (Figure S8C). LcrV, which did not activate NLRC4 (Figure S8B), shares a relatively high level (36.3%) of identity with PcrV but differs from PcrV in the T6R (Figure S8D). Substitution of the T6R of LcrV with the T5R of PcrV enabled the mutant, LcrV-T6RM, to gain the ability to activate NLRC4 inflammasome (Figure S8D). Similarly, substitution of the T5R of EspA_EHEC_ with the T6R of EseB enabled the mutant, EspA_EHEC_-T5RM, to activate NLRC4 inflammasome (Figure S8E).

**Figure 5.**
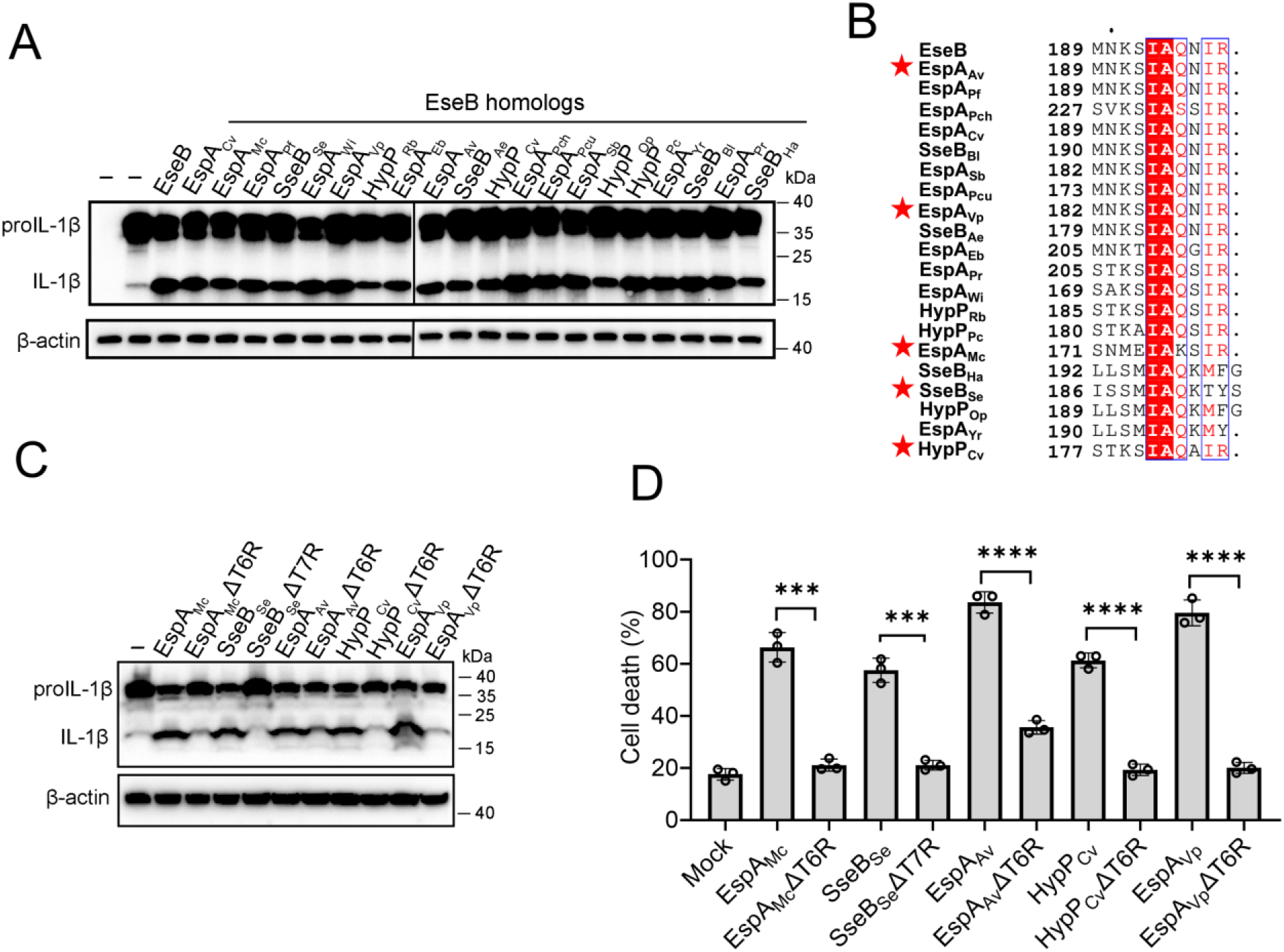
The ability of the EseB homologs to activate the NLRC4/NAIP inflammasome. (**A**) NLRC4 inflammasome-reconstituted HEK293T cells were transfected with or without EseB homologs (Table S2). The cells were immunoblotted with antibodies against IL-1β and β-actin (loading control). (**B**) Sequence alignment of the C-terminal regions of the EseB homologs. Red stars indicate the EseB selected for mutation analysis. (**C**) NLRC4 inflammasome-reconstituted HEK293T cells expressing or not expressing the indicated EseB homologs or their mutants were immunoblotted as panel A. (**D**) THP-1 cells were electroporated with the indicated EseB homologs or their mutants or PBS (mock) and then determined for cell death. Data are the means of triplicate assays and shown as means ± SD. ns, not significant, \*\*\**p*<0.001, \*\*\*\**p*<0.0001, Student’s *t*-test.

## Discussion

In this study, we examined the mechanism of inflammasome-mediated pyroptosis induced by bacterial T3SS translocon. Well-known inflammasome proteins, such as NLRP3 and NLRC4, and Casp4 are intracellular PRRs that induce pyroptosis during intracellular bacterial infections [27, 28]. While NLRP3 activation can occur upon alterations in cellular homeostasis [29-31], NLRC4 and Casp4 are primarily activated by specific PAMP ligands presented by microbial organisms [1, 32]. Previous studies have demonstrated that *E. tarda* components or secreted particles can cause pyroptosis in murine macrophages and human epithelial cells [33-37]. In this study, we found that in the infection model of THP-1 derived human macrophages, which express multiple inflammasomes, *E. tarda* induced pyroptosis in a manner that depended on NLRC4, NLRP3, ASC, Casp1, and Casp4. This observation indicated that *E. tarda* infection activated both the canonical and the non-canonical inflammasome pathways, which might be due to the multiplicity of virulence factors expressed by *E. tarda*. In line with this result, cell death was nearly blocked when the bacteria were prevented from entering the host cell cytoplasm, suggesting that *E. tarda*-triggered cell death was an event that required direct interaction between the bacteria and the inflammasome molecules.

T3SS is one of the critical armaments of intracellular bacteria to combat the host’s defense system [17, 38]. T3SS delivers multiple bacterial effectors into host cells, which manipulate host immune responses to foster bacterial survival and expansion [39, 40]. This feature makes the T3SS apparatus readily exposed to the host cytosol, thus susceptible to detection by host receptors such as inflammasomes [6-9, 40]. Studies have demonstrated that in mice, T3SS needle protein, T3SS rod protein, and flagellin are directly detected by NAIP1, NAIP2, and NAIP5/6, respectively [7, 9]. In contrast, in humans, T3SS needle/rod proteins and flagellin are all detected by a single NAIP [8, 10, 41]. For *E. tarda,* two reports showed that flagellin was required to induce fish macrophage death [33] but not required to induce murine macrophage death [35]. In our study, we found that flagellin was dispensable for *E. tarda*-induced pyroptosis of human macrophages. Consistently, the flagellin proteins were unable to activate human NAIP/NLRC4 inflammasomes. This observation differs from that of other pathogen flagellin proteins, which are recognized by NLRC4 inflammasomes [7-8, 41]. Like flagellin, the *E. tarda* rod protein EsaI also failed to activate the NLRC4 inflammasome. These results indicate that *E. tarda* flagellin and rod evade recognition by human NLRC4, probably as a strategy to facilitate bacterial infection. Unlike the needle, rod proteins and flagellin, T3SS translocon proteins have seldom been reported to promote inflammasome activation and pyroptosis. Limited studies showed that the *Yersinia* translocon proteins YopD and YopB could translocate into host cells and indirectly activate inflammasome, resulting in cell death [42-44]; the translocon proteins of *Pseudomonas aeruginosa* PopB–PopD may be associated with inflammasome activation [45]. Currently, it is unknown whether any inflammasome can directly recognize the translocon proteins. In the present study, mutational analyses showed that the translocon proteins were essential for *E. tarda* to induce pyroptosis and activate the non-canonical inflammasome Casp4. In particular, we found that the translocon protein EseB, when present intracellularly, sufficed to cause pyroptosis via NLRC4/NAIP. Moreover, the terminal residues proved to be vital for EseB activity, and the CT region alone could trigger pyroptosis in a manner similar to EseB. Consistently, although EseB differs dramatically from the NLRC4/NAIP-stimulating needle proteins in the NT region, it shares notable identities with the needle proteins in the CT region, suggesting a conserved mechanism of inflammasome activation via the CT region in these proteins.

Although T3SS is present in many bacterial pathogens, the particular mechanisms of T3SS translocon proteins, notably EseB, to induce host immune response are unclear. In this study, we identified EseB homologs in a large number of bacteria with T3SS and found that, like *E. tarda* EseB, all of these identified EseB homologs were able to activate NLRC4/NAIP, thus suggesting the wide existence of bacteria−host interaction mediated by EseB and the NLRC4/NAIP inflammasome. Sequence analysis revealed highly conserved T6R in all of the EseB homologs, which was crucial to NLRC4 inflammasome activation. Similar observation has been reported for the needle protein of T3SS. It has been shown that deletion of the last five amino acids from the needle protein prevented its self-association, so that the protein could exist only in the monomeric form, indicating the importance of these terminal amino acids in maintaining the stability of the high order structure of the protein [46-48]. In support of this, the cryo-electron microscopic structure of the needle−HsNAIP−HsNLRC4 complex showed that the last few amino acids of the needle were involved in interaction with human NAIP [49]. The importance of the last few residues is also suggested by the report that *S. typhimurium* SPI2 T3SS rod protein SsaI may evade NLRC4/NAIP inflammasome recognition by alteration in the last 8 amino acids [6]. In our study, we found that replacement of the terminal residues of EspA_EHEC_ with that of EseB switched EspA_EHEC_ to an NLRC4 activator. Similar observations were made with LcrV. These results suggest that the lack of EseB T6R-like terminal residues might be a strategy for EspA_EHEC_ and LcrV to evade host immune detection. However, we also observed that IpaC and SipC, which possess EseB T6R-like terminal residues, failed to activate the NLRC4 inflammasome. Together these observations indicate that the terminal amino acids are a key, but not the sole, determinant in NLRC4 activation. Future studies are needed to find out the additional determinant(s) that works with the terminal residues to activate the NLRC4-mediated signaling.

In summary, our study demonstrated the importance and the mechanism of bacterial T3SS translocon in host interaction. We found that the translocon protein, EseB, triggered pyroptosis by directly activating the host NLRC4/NAIP inflammasome via the CT region, especially the T6R. Both the sequence and the inflammasome-stimulating function of the T6R were highly conserved in the EseB homologs of diverse bacteria. However, it must be said that for a translocator protein, the possession of a conserved T6R alone does not suffice to activate the NLRC4 inflammasome. These findings deepened the understanding of the function and mechanism of T3SS-mediated interaction between pathogens and hosts.

## Acknowledgements

This work was supported by the Science & Technology Innovation Project of Laoshan Laboratory (LSKJ202203000), the National Natural Science Foundation of China (31330081), and the National Key Research and Development Project of China (2018YFD0900500).

## Conflict of Interests

The authors declare that they have no conflict of interest.

## Author contributions

L.S. and Y.Z. conceived the study and wrote the paper; L.S. obtained the funding; Y.Z. and H.S.Z. conducted the experiments; Y.Z., H.S.Z., J.Q.L and X.H analyzed the data.

## Notes

### Competing Interest Statement

The authors have declared no competing interest.

### Summary of Updates

Section on Results and Discussion updated; Figure 5 revised; author affiliations updated; Supplemental files updated.

## References

1. Christgen S, Place DE, Kanneganti TD. Toward targeting inflammasomes: insights into their regulation and activation. Cell Res. 2020;30(4):315–27. doi: 10.1038/s41422-020-0295-8.

2. Kanneganti TD. Intracellular innate immune receptors: Life inside the cell. Immunol Rev. 2020;297(1):5–12. doi: 10.1111/imr.12912.

3. He W, Wan H, Hu L, Chen P, Wang X, Huang Z, et al. Gasdermin D is an executor of pyroptosis and required for interleukin-1β secretion. Cell Res. 2015;25(12):1285–98. doi: 10.1038/cr.2015.139.

4. Shi J, Zhao Y, Wang K, Shi X, Wang Y, Huang H, et al. Cleavage of GSDMD by inflammatory caspases determines pyroptotic cell death. Nature. 2015;526(7575):660-5. doi: 10.1038/nature15514.

5. Kayagaki N, Stowe IB, Lee BL, O’Rourke K, Anderson K, Warming S, et al. Caspase-11 cleaves gasdermin D for non-canonical inflammasome signalling. Nature. 2015;526(7575):666-71. doi: 10.1038/nature15541.

6. Miao EA, Mao DP, Yudkovsky N, Bonneau R, Lorang CG, Warren SE, et al. Innate immune detection of the type III secretion apparatus through the NLRC4 inflammasome. Proc Natl Acad Sci U S A. 2010;107(7):3076–80. doi: 10.1073/pnas.0913087107.

7. Zhao Y, Yang J, Shi J, Gong YN, Lu Q, Xu H, et al. The NLRC4 inflammasome receptors for bacterial flagellin and type III secretion apparatus. Nature. 2011;477(7366):596-600. doi: 10.1038/nature10510.

8. Reyes Ruiz VM, Ramirez J, Naseer N, Palacio NM, Siddarthan IJ, Yan BM, et al. Broad detection of bacterial type III secretion system and flagellin proteins by the human NAIP/NLRC4 inflammasome. Proc Natl Acad Sci U S A. 2017;114(50):13242–7. doi: 10.1073/pnas.1710433114.

9. Kofoed EM, Vance RE. Innate immune recognition of bacterial ligands by NAIPs determines inflammasome specificity. Nature. 2011;477(7366):592-5. doi: 10.1038/nature10394.

10. Yang J, Zhao Y, Shi J, Shao F. Human NAIP and mouse NAIP1 recognize bacterial type III secretion needle protein for inflammasome activation. Proc Natl Acad Sci U S A. 2013;110(35):14408–13. doi: 10.1073/pnas.1306376110.

11. Sui ZH, Xu HJ, Wang HD, Jiang S, Chi H, Sun L. Intracellular trafficking pathways of *Edwardsiella tarda*: from clathrin-and caveolin-mediated endocytosis to endosome and lysosome. Front Cell Infect Microbiol. 2017;7. doi: 10.3389/fcimb.2017.00400.

12. Li HL, Sun BG, Ning XH, Jiang S, Sun L. A comparative analysis of *Edwardsiella tarda*-induced transcriptome profiles in RAW264. 7 cells reveals new insights into the strategy of bacterial immune evasion. Int J Mol Sci. 2019;20(22). doi: 10.3390/ijms20225724.

13. Leung KY, Siame BA, Tenkink BJ, Noort RJ, Mok YK. *Edwardsiella tarda*-virulence mechanisms of an emerging gastroenteritis pathogen. Microbes Infect. 2012;14(1):26–34. doi: 10.1016/j.micinf.2011.08.005.

14. Leung KY, Wang Q, Yang Z, Siame BA. *Edwardsiella piscicida*: A versatile emerging pathogen of fish. Virulence. 2019;10(1):555–67. doi: 10.1080/21505594.2019.1621648.

15. Hirai Y, Asahata-Tago S, Ainoda Y, Fujita T, Kikuchi K. *Edwardsiella tarda* bacteremia. A rare but fatal water-and foodborne infection: Review of the literature and clinical cases from a single centre. Can J Infect Dis Med Microbiol. 2015;26(6):313–8. doi: 10.1155/2015/702615.

16. Leung KY, Wang Q, Zheng X, Zhuang M, Yang Z, Shao S, et al. Versatile lifestyles of *Edwardsiella*: Free-living, pathogen, and core bacterium of the aquatic resistome. Virulence. 2022;13(1):5–18. doi: 10.1080/21505594.2021.2006890.

17. Portaliou AG, Tsolis KC, Loos MS, Zorzini V, Economou A. Type III secretion: building and operating a remarkable nanomachine. Trends Biochem Sci. 2016;41(2):175–89. doi: 10.1016/j.tibs.2015.09.005.

18. Dey S, Chakravarty A, Biswas PG, De Guzman RN. The type III secretion system needle, tip, and translocon. Protein Sci. 2019;28(9):1582–93. doi: 10.1002/pro.3682.

19. Tan YP, Zheng J, Tung SL, Rosenshine I, Leung KY. Role of type III secretion in *Edwardsiella tarda* virulence. Microbiology (Reading). 2005;151(7):2301–13. doi: 10.1099/mic.0.28005-0.

20. Wang B, Mo ZL, Mao YX, Zou YX, Xiao P, Li J, et al. Investigation of EscA as a chaperone for the *Edwardsiella tarda* type III secretion system putative translocon component EseC. Microbiology (Reading). 2009;155(4):1260–71. doi: 10.1099/mic.0.021865-0.

21. Okuda J, Kiriyama M, Suzaki E, Kataoka K, Nishibuchi M, Nakai T. Characterization of proteins secreted from a type III secretion system of *Edwardsiella tarda* and their roles in macrophage infection. Dis Aquat Organ. 2009;84(2):115–21. doi: 10.3354/dao02033.

22. Zhao Y, Jiang S, Zhang J, Guan XL, Sun BG, Sun L. A virulent *Bacillus cereus* strain from deep-sea cold seep induces pyroptosis in a manner that involves NLRP3 inflammasome, JNK pathway, and lysosomal rupture. Virulence. 2021;12(1):1362–76. doi: 10.1080/21505594.2021.1926649.

23. Wang Y, Luo J, Zhao Y, Zhang J, Guan X, Sun L. Haemolysins are essential to the pathogenicity of deep-sea *Vibrio fluvialis*. iScience. 2024;27(5). doi: 10.1016/j.isci.2024.109558.

24. Li M, Wu M, Sun Y, Sun L. *Edwardsiella tarda* TraT is an anti-complement factor and a cellular infection promoter. Commun Biol. 2022;5(1):637. doi: 10.1038/s42003-022-03587-3.

25. Liu X, Wang X, Sun B, Sun L. The involvement of thiamine uptake in the virulence of *Edwardsiella piscicida*. Pathogens. 2022;11(4). doi: 10.3390/pathogens11040464.

26. Zhao Y, Sun L. *Bacillus cereus* cytotoxin K triggers gasdermin D-dependent pyroptosis. Cell Death Discov. 2022;8(1):305. doi: 10.1038/s41420-022-01091-5.

27. Storek KM, Monack DM. Bacterial recognition pathways that lead to inflammasome activation. Immunol Rev. 2015;265(1):112–29. doi: 10.1111/imr.12289.

28. Broz P, Dixit VM. Inflammasomes: mechanism of assembly, regulation and signalling. Nat Rev Immunol. 2016;16(7):407–20. doi: 10.1038/nri.2016.58.

29. Gong T, Yang Y, Jin T, Jiang W, Zhou R. Orchestration of NLRP3 inflammasome activation by ion fluxes. Trends Immunol. 2018;39(5):393–406. doi: 10.1016/j.it.2018.01.009.

30. Hughes MM, O’Neill LA. Metabolic regulation of NLRP3. Immunol Rev. 2018;281(1):88–98. doi: 10.1111/imr.12608.

31. Seoane PI, Lee B, Hoyle C, Yu S, Lopez-Castejon G, Lowe M, et al. The NLRP3–inflammasome as a sensor of organelle dysfunction. J Cell Biol. 2020;219(12):e202006194. doi: 10.1083/jcb.202006194.

32. Egan MS, Zhang J, Shin S. Human and mouse NAIP/NLRC4 inflammasome responses to bacterial infection. Curr Opin Microbiol. 2023;73:102298. doi: 10.1016/j.mib.2023.102298.

33. Xie HX, Lu JF, Rolhion N, Holden DW, Nie P, Zhou Y, et al. *Edwardsiella tarda*-induced cytotoxicity depends on its type III secretion system and flagellin. Infect Immun. 2014;82(8):3436–45. doi: 10.1128/IAI.01065-13.

34. Zhang L, Ni C, Xu W, Dai T, Yang D, Wang Q, et al. Intramacrophage infection reinforces the virulence of *Edwardsiella tarda*. J Bacteriol. 2016;198(10):1534–42. doi: 10.1128/JB.00978-15.

35. Chen H, Yang D, Han F, Tan J, Zhang L, Xiao J, et al. The bacterial T6SS effector EvpP prevents NLRP3 inflammasome activation by inhibiting the Ca^2+^-dependent MAPK-JNK pathway. Cell Host Microbe. 2017;21(1):47–58. doi: 10.1016/j.chom.2016.12.004.

36. Chen S, Yang D, Wen Y, Jiang Z, Zhang L, Jiang J, et al. Dysregulated hemolysin liberates bacterial outer membrane vesicles for cytosolic lipopolysaccharide sensing. PLoS Pathog. 2018;14(8):e1007240. doi: 10.1371/journal.ppat.1007240.

37. Xu W, Gu Z, Zhang L, Zhang Y, Liu Q, Yang D. *Edwardsiella piscicida* virulence effector trxlp promotes the NLRC4 inflammasome activation during infection. Microb Pathog. 2018;123:496–504. doi: 10.1016/j.micpath.2018.08.016.

38. Deng W, Marshall NC, Rowland JL, McCoy JM, Worrall LJ, Santos AS, et al. Assembly, structure, function and regulation of type III secretion systems. Nat Rev Microbiol. 2017;15(6):323–37. doi: 10.1038/nrmicro.2017.20.

39. Raymond B, Young JC, Pallett M, Endres RG, Clements A, Frankel G. Subversion of trafficking, apoptosis, and innate immunity by type III secretion system effectors. Trends Microbiol. 2013;21(8):430–41. doi: 10.1016/j.tim.2013.06.008.

40. Ratner D, Orning MP, Lien E. Bacterial secretion systems and regulation of inflammasome activation. J Leukoc Biol. 2017;101(1):165–81. doi: 10.1189/jlb.4MR0716-330R.

41. Kortmann J, Brubaker SW, Monack DM. Cutting edge: inflammasome activation in primary human macrophages is dependent on flagellin. J Immunol. 2015;195(3):815–9. doi: 10.4049/jimmunol.1403100.

42. Casson CN, Copenhaver AM, Zwack EE, Nguyen HT, Strowig T, Javdan B, et al. Caspase-11 activation in response to bacterial secretion systems that access the host cytosol. PLoS Pathog. 2013;9(6):e1003400. doi: 10.1371/journal.ppat.1003400.

43. Zwack EE, Snyder AG, Wynosky-Dolfi MA, Ruthel G, Philip NH, Marketon MM, et al. Inflammasome activation in response to the *Yersinia* type III secretion system requires hyperinjection of translocon proteins YopB and YopD. mBio. 2015;6(1):e02095–14. doi: 10.1128/mBio.02095-14.

44. Zwack EE, Feeley EM, Burton AR, Hu B, Yamamoto M, Kanneganti TD, et al. Guanylate binding proteins regulate inflammasome activation in response to hyperinjected *Yersinia* translocon components. Infect Immun. 2017;85(10). doi: 10.1128/IAI.00778-16.

45. Dortet L, Lombardi C, Cretin F, Dessen A, Filloux A. Pore-forming activity of the *Pseudomonas aeruginosa* type III secretion system translocon alters the host epigenome. Nat Microbiol. 2018;3(3):378–86. doi: 10.1038/s41564-018-0109-7.

46. Deane JE, Roversi P, Cordes FS, Johnson S, Kenjale R, Daniell S, et al. Molecular model of a type III secretion system needle: Implications for host-cell sensing. Proc Natl Acad Sci U S A. 2006;103(33):12529–33. doi: 10.1073/pnas.0602689103.

47. Zhang L, Wang Y, Picking WL, Picking WD, De Guzman RN. Solution structure of monomeric BsaL, the type III secretion needle protein of *Burkholderia pseudomallei*. J Mol Biol. 2006;359(2):322–30. doi: 10.1016/j.jmb.2006.03.028.

48. Wang Y, Ouellette AN, Egan CW, Rathinavelan T, Im W, De Guzman RN. Differences in the electrostatic surfaces of the type III secretion needle proteins PrgI, BsaL, and MxiH. J Mol Biol. 2007;371(5):1304–14. doi: 10.1016/j.jmb.2007.06.034.

49. Matico RE, Yu X, Miller R, Somani S, Ricketts MD, Kumar N, et al. Structural basis of the human NAIP/NLRC4 inflammasome assembly and pathogen sensing. Nat Struct Mol Biol. 2024;31(1):82–91. doi: 10.1038/s41594-023-01143-z.

