## Supplemental data for "T3SS translocon induces pyroptosis by direct interaction with NLRC4/NAIP inflammasome"

**Table S1.** The primers used for the construction of *Edwardsiella tarda* translocon mutants.

| Primer | Sequence (5’-3’)[a] | Mutant |
|---|---|---|
| EseB/D-F1 | GGTTACCCGCATGCAGCAGCGCGCAGGCAAACGCG | |
| EseB/D R1 | AGAGCACTAGGAGGCACGATGGATGACG | Δ*eseB-D* |
| EseB/D-F2 | TGCCTCCTAGTGCTCTCCTCTGAGGGAT | |
| EseB/D-R2 | CCCTTCTAGATAGATCTGGAGACGCCGCTCAACGCCT | |
| EscA-F1 | GGTTACCCGCATGCACTGCGCCTTAGCCTGATCCTC | |
| EscA-R1 | CGCTGCATCGTCAGTTCACACCGGTGACC | Δ*escA* |
| EscA-F2 | AACTGACGATGCAGCGTAGCGAGATCGTC | |
| EscA-R2 | CCCTTCTAGATAGATCTCCAGTTCATGACGCTATTCAC | |
| EseB-F1 | GGTTACCCGCATGCAGTCTCCTCGGTCACCGGTGTG | |
| EseB-R1 | TAAACCACGCCGAGATGAACAAATCCATC | Δ*eseB* |
| EseB-F2 | ATCTCGGCGTGGTTTACACCGCCGTGGTA | |
| EseB-R2 | CCCTTCTAGATAGATCTGATAACCAAACAGCAAACGTT | |
| EseC-F1 | GGTTACCCGCATGCACCGCTTAAGCTGCTGCTCGGC | |
| EseC-R1 | CTGAGACCGGCGCCATGCTCAGCAACATC | Δ*eseC* |
| EseC-F2 | ATGGCGCCGGTCTCAGTGATATTGTTCAT | |
| EseC-R2 | CCCTTCTAGATAGATCTAACAAATCCATCGCCCAGAAT | |
| EseD-F1 | GGTTACCCGCATGCATTATCAGGGCGTGCGCCCCGG | |
| EseD-R1 | ACGGTGTTGCCCACGAACGTATCGCCAGC | Δ*eseD* |
| EseD-F2 | TCGTGGGCAACACCGTGGCTGCCGCTGTC | |
| EseD-R2 | CCCTTCTAGATAGATCTGCCGAGCAGAGCGGTCGCTTC | |
| FliC-F1 | GGTTACCCGCATGCAGTATACAAATCAGTCACGTCG | |
| FliC-R1 | CGCTTCACCGTACAGAACCGTTTCGATTCC | Δ*fliC* |
| FliC-F2 | GTTCTGTACGGTGAAGCGGTTGGAGATCG | |
| FliC-R2 | CCCTTCTAGATAGATCTTACGCGTTATCGGCTCTGTTG | |

[a]The underlined sequences are the homologous arm sequences in pDM4.

**Table S2.** Translocator proteins used in this study

| Protein | Bacteria | Accession number* | Identity (%) with *Edwardsiella tarda* EseB |
|---|---|---|---|
| EspA$_{Av}$ | *Aeromonas veronii* | QMS76431 | 77.3 |
| EspA$_{Pf}$ | *Pseudogulbenkiania ferrooxidans* | ERE19475 | 76.8 |
| SseB$_{Bl}$ | *Burkholderia lata* | VWD65036 | 64.2 |
| EspA$_{Sb}$ | *Shewanella baltica* | AEH13802 | 51.0 |
| EspA$_{Pcu}$ | *Parashewanella curva* | RLV59315 | 50.5 |
| SseB$_{Ae}$ | *Arsenophonus endosymbiont* | A0A3B0MKK6 (UniProtKB accession number) | 50.2 |
| EspA$_{Vp}$ | *Vibrio pectenicida* | NOH71600 | 48.8 |
| EspA$_{Eb}$ | *Enterobacteriaceae bacterium* | QLK63763 | 37.7 |
| HypP$_{Rb}$ | *Rouxiella badensis* | ORJ26452 | 36.5 |
| EspA$_{Pr}$ | *Providencia rettgeri* | AVL73471 | 35.4 |
| EspA$_{Wi}$ | *Winslowiella iniecta* | KOC86666 | 34.8 |
| EspA$_{Mc}$ | *Mycoavidus cysteinexigens* | BBE08751 | 34.1 |
| EspA$_{Cv}$ | *Chromobacterium vaccinii* | AVG16459 | 33.3 |
| HypP$_{Cv}$ | *Chromobacterium violaceum* | AAQ60250 | 33.3 |
| SseB$_{Se}$ | *Salmonella enterica* SPI-2 | HAF8569287 | 33.0 |
| HypP$_{Pc}$ | *Pantoea cypripedii* | QGY32172 | 31.1 |
| SseB$_{Ha}$ | *Hafnia alvei* | SCM50665 | 30.6 |
| EspA$_{Yr}$ | *Yokenella regensburgei* | QIU89219 | 30.0 |
| HypP$_{Op}$ | *Obesumbacterium proteus* | AMO83764 | 29.7 |
| EspA$_{Pch}$ | *Pseudomonas chlororaphis* | QLL16699 | 28.5 |
| EspA$_{EPEC}$ | enteropathogenic *Escherichia coli* | WP_000381567 | 22.8 |
| EspA$_{EHEC}$ | enterohaemorrhagic *Escherichia coli* | WP_000381516 | 20.4 |
| EspA$_{Ea}$ | *Escherichia albertii* | WP_000381555 | 20.4 |
| EspA$_{Cr}$ | *Citrobacter rodentium* | AAL06381 | 18.8 |
| IpaC | *Shigella flexneri* | NP_858260 | 13.0 |
| IpaD | *Shigella flexneri* | ADA76864 | 12.9 |
| SipD | *Salmonella enterica* SPI-1 | NP_461804 | 12.8 |
| SipC | *Salmonella enterica* SPI-1 | NP_461805 | 10.4 |
| YopD | *Yersinia enterocolitica* | WP_010891207 | 9.7 |
| PcrV | *Pseudomonas aeruginosa* | NP_250397 | 9.6 |
| LcrV | *Yersinia enterocolitica* | WP_014609483 | 9.1 |
| PopD | *Pseudomonas aeruginosa* | NP_250400 | 8.0 |
| SseD | *Salmonella enterica* SPI-2 | AAC28882 | 7.4 |

| | | | |
|---|---|---|---|
| EspB | *enterohaemorrhagic Escherichia coli* | NP_312581 | 4.0 |

*, GenBank accession number unless otherwise indicated.

HypP, hypothetical or unnamed protein

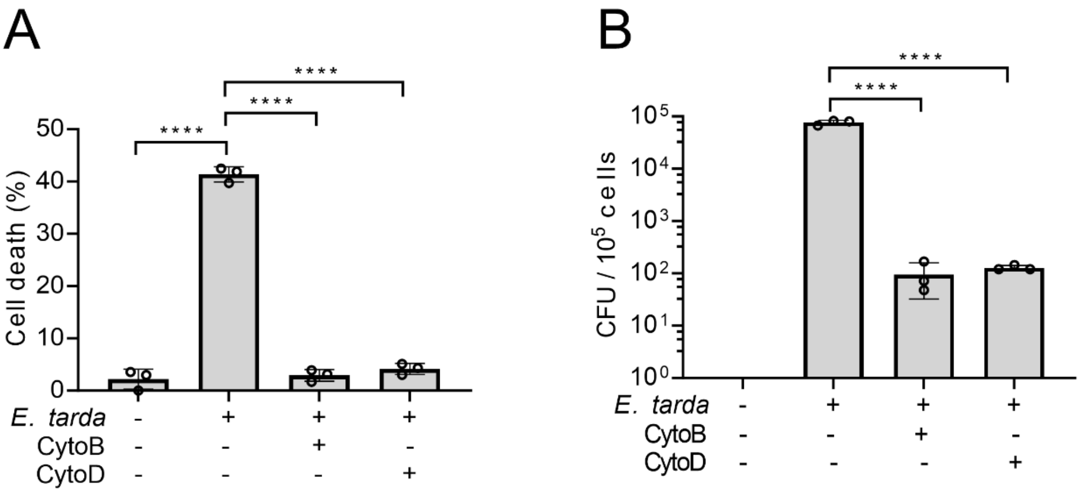


**Figure S1. The effect of cytochalasin B (CytoB) and cytochalasin D (CytoD) on the ability of *Edwardsiella tarda* to induce cell death.** (**A**) dTHP-1 cells pretreated with or without DMSO, CytoB, or CytoD were infected with or without *E. tarda* for 2 h and then determined for cell death. (**B**) dTHP-1 cells pretreated with or without DMSO, CytoB, or CytoD were infected with or without *E. tarda* for 1 h. The extracellular bacteria were killed by gentamycin. The intracellular bacteria were quantified by plate count. Data are the means of triplicate assays and shown as means ± SD. ****$p$<0.0001, one-way ANOVA with Dunnett's multiple-comparison test.

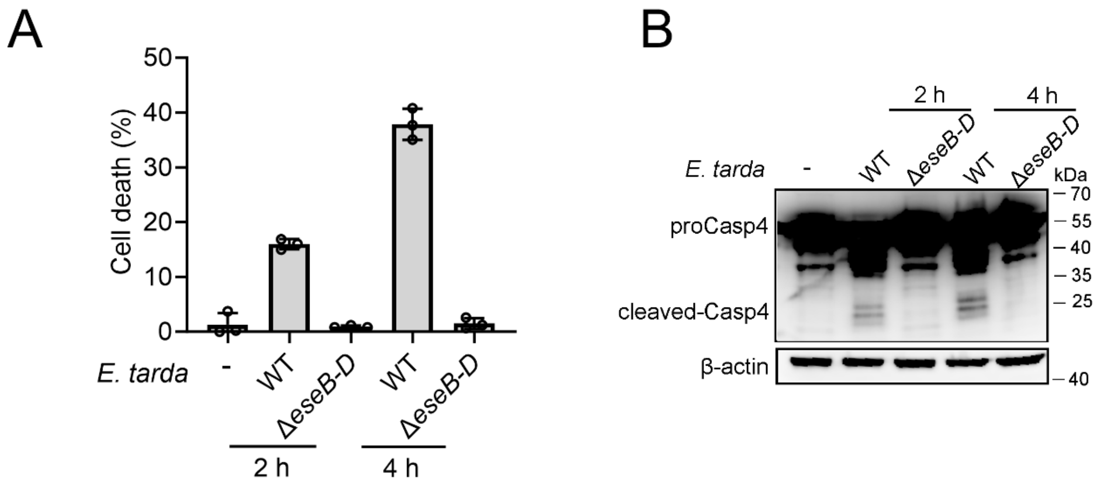


**Figure S2. The effect of Δ*eseB-D* on Casp4 activation.** (A) dTHP-1 cells were treated with wild type (WT) *Edwardsiella tarda* or the Δ*eseB-D* mutant for 2 h or 4 h. The cells were subjected to cell death analysis (A) and immunoblot with antibodies against Casp4 and β-actin (B). Data in panel A are the means of triplicate assays and shown as means ± SD.

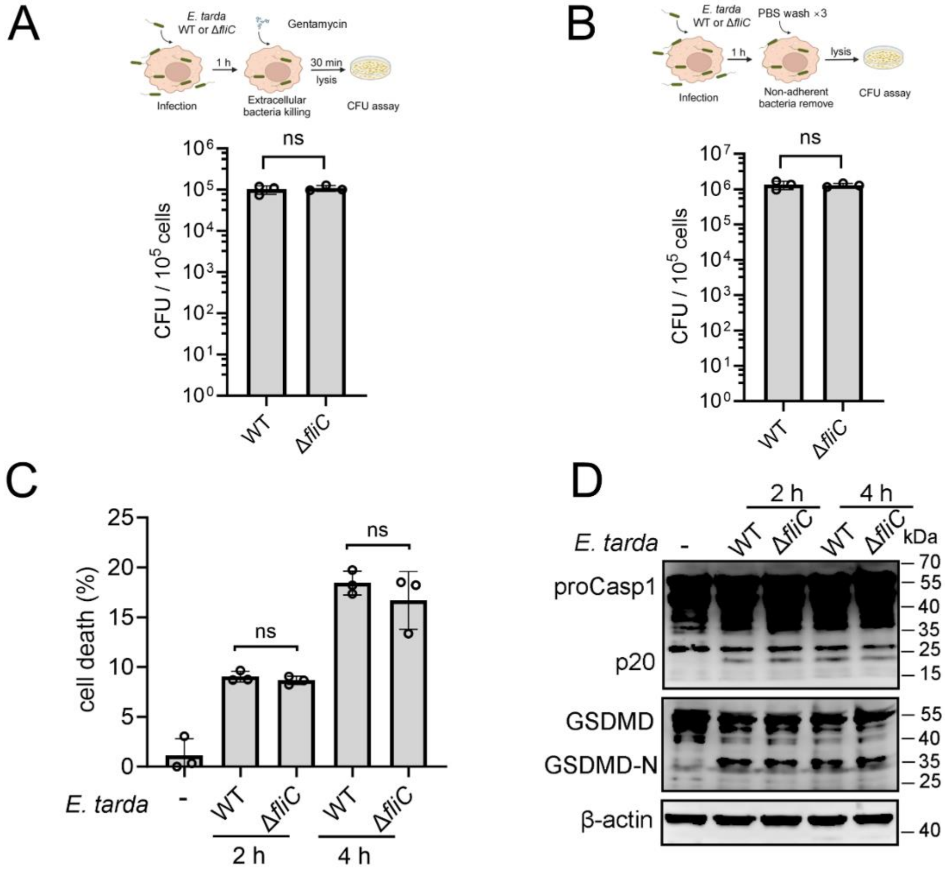


**Figure S3. The involvement of flagellin in *Edwardsiella tarda*-induced pyroptosis of human macrophages.** (A, B) dTHP-1 cells were infected with *E. tarda* wild type (WT) or Δ*fliC* for 1h. The intracellular bacteria (A) and the total bacteria associated with the cells (B) were determined by plate count. (C, D) dTHP-1 cells were treated with or without *E. tarda* WT or Δ*fliC* for 2 or 4 h, and then subjected to cell death analysis (C) and immunoblotting with antibodies against Casp1, GSDMD, and β-actin (D). Data in panels A-C are the means of triplicate assays and shown as means ± SD. ns, not significant; $p>0.05$, Student's *t*-test.

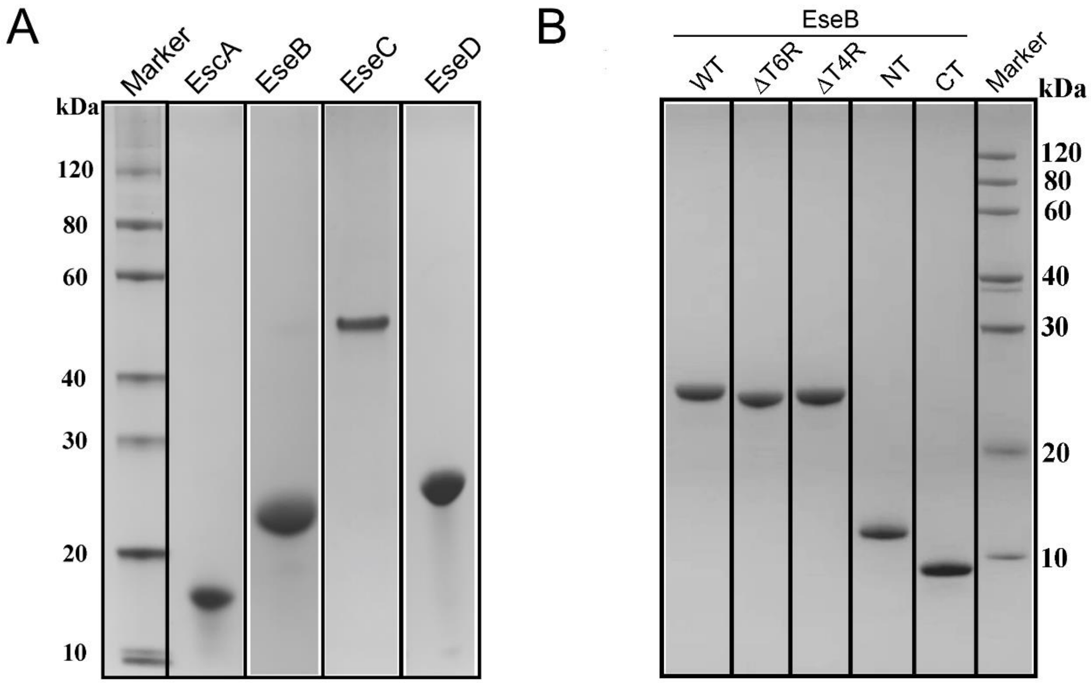


**Figure S4. SDS-PAGE analysis of purified recombinant proteins.** Purified recombinant translocon proteins (EscA, EseB, EseC, and EseD) (A) and EseB truncates (B) were subjected to SDS-PAGE and stained with Coomassie brilliant blue R-250. WT, wild type.

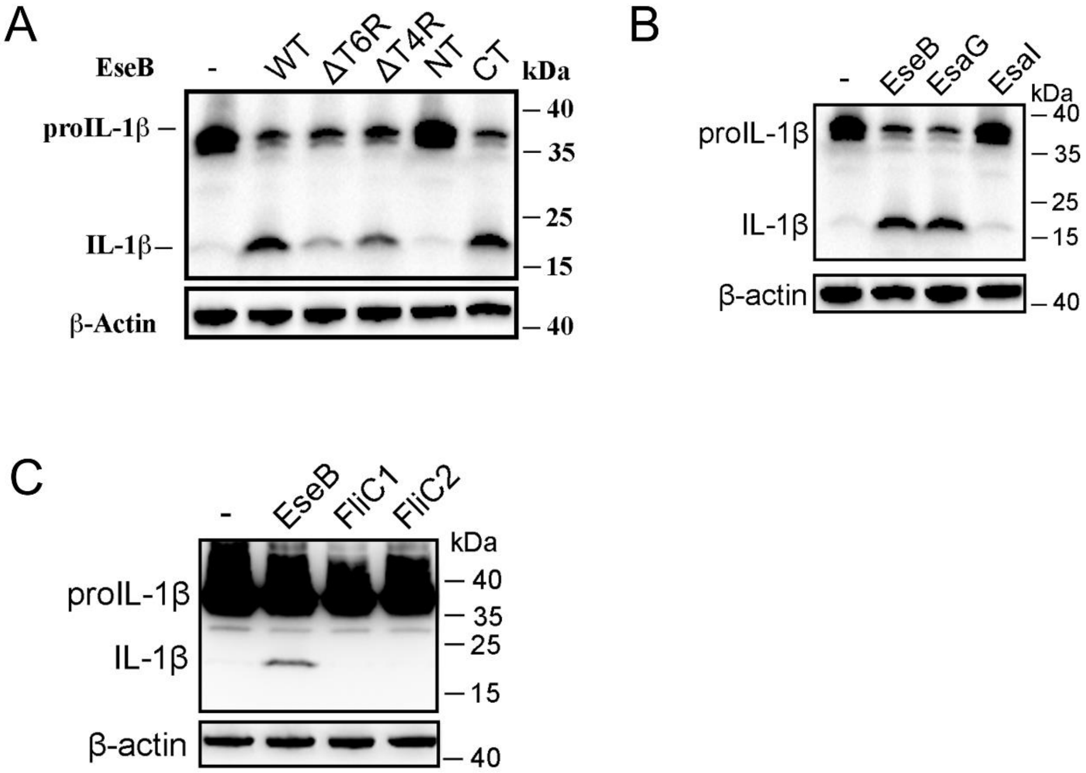


**Figure S5. The ability of *Edwardsiella tarda* EseB, rod (EsaI), needle (EsaG), and flagellin (FliC1/2) to activate NLRC4.** NLRC4 inflammasome-reconstituted HEK293T cells expressing or not expressing wild type (WT) or mutant EseB (**A**), EsaI/EsaG (**B**), or FliC1/2 (**C**) for 24 h were immunoblotted with antibodies against IL-1β and β-actin (loading control).

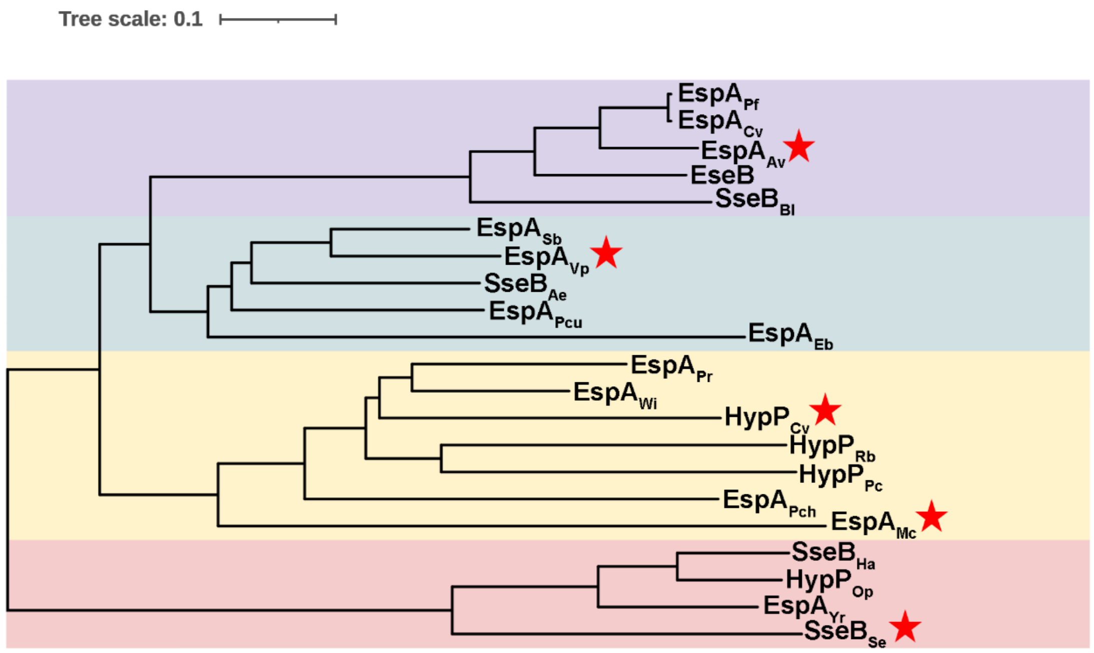


**Figure S6. Phylogenetic analysis of EseB homologs**. The phylogenetic tree was generated using ClustalW alignment and the Neighbor Joining method. Red pentagrams indicate the bacteria whose EseB homologs were selected for mutation analysis.

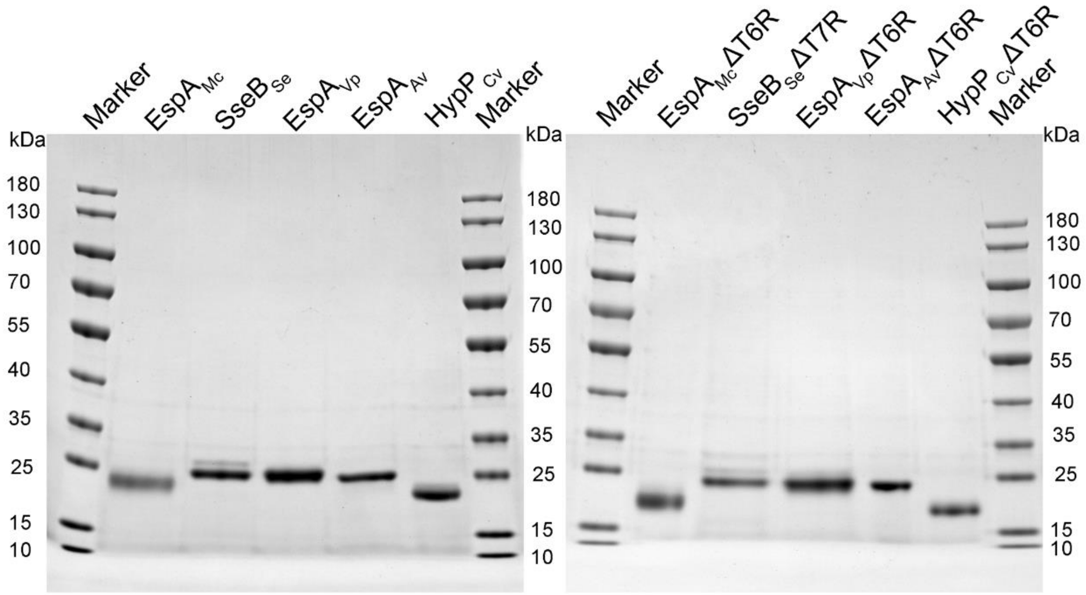


**Figure S7. SDS-PAGE analysis of purified recombinant proteins.** Purified recombinant EseB homologues (left) and their mutants (right) were subjected to SDS-PAGE and stained with Coomassie brilliant blue R-250. The full names of the proteins are shown in Table S2.

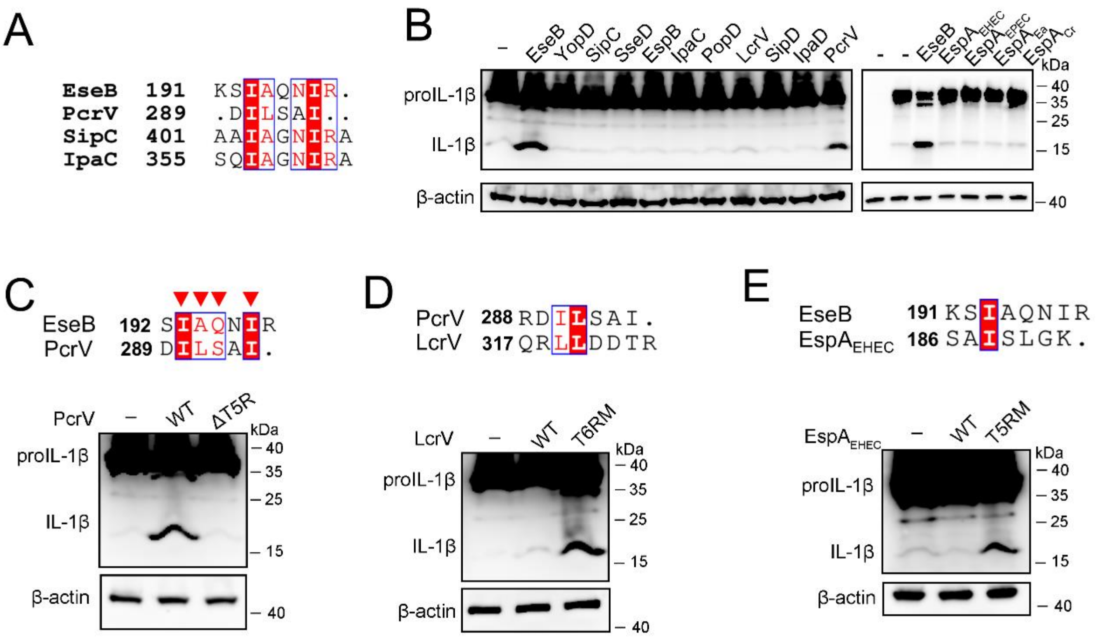


**Figure S8. The effects of translocator proteins on NLRC4/NAIP inflammasome activation. (A)** Sequence alignment of the C-terminal regions of EseB, PcrV, SipC and IpaC. (**B**) NLRC4 inflammasome-reconstituted HEK293T cells expressing or not expression (-) the indicated proteins for 24 h were immunoblotted with antibodies against IL-1β and β-actin (loading control). (**C-E**) NLRC4 inflammasome-reconstituted HEK293T cells expressing or not expression (-) wild type (WT) and mutant PcrV (C), LcrV (D), and $\mathrm{EspA_{EHEC}}$ (E) were immunoblotted as above. The C-terminal regions of PcrV, LcrV, and $\mathrm{EspA_{EHEC}}$ were aligned on top of the panels.
